# Signal propagation via cortical hierarchies

**DOI:** 10.1101/2020.02.15.950907

**Authors:** Bertha Vázquez-Rodríguez, Zhen-Qi Liu, Patric Hagmann, Bratislav Mišić

**Author notes:** These authors contributed equally to this work (Bertha Vázquez-Rodríguez, Zhen-Qi Liu).

## Abstract

The wiring of the brain is organized around a putative unimodal-transmodal hierarchy. Here we investigate how this intrinsic hierarchical organization of the brain shapes the transmission of information among regions. The hierarchical positioning of individual regions was quantified by applying diffusion map embedding to resting state functional MRI networks. Structural networks were reconstructed from diffusion spectrum imaging and topological shortest paths among all brain regions were computed. Sequences of nodes encountered along a path were labelled by their hierarchical position, tracing out path motifs. We find that the cortical hierarchy guides communication in the network. Specifically, nodes are more likely to forward signals to nodes closer in the hierarchy and cover a range of unimodal and transmodal regions, potentially enriching or diversifying signals en route. We also find evidence of systematic detours, particularly in attention networks, where communication is re-routed. Altogether, the present work highlights how the cortical hierarchy shapes signal exchange and imparts behaviourally-relevant communication patterns in brain networks.

## INTRODUCTION

Adaptive behaviour requires transmission of information between neuronal populations. The architecture of white matter networks supports an array of signal propagation patterns, linking sensation, cognition and action [3]. Brain networks, reconstructed from multiple species and at multiple spatial scales, possess multiple nonrandom attributes that make such flexible communication possible, including near-minimal path length and high clustering [28, 32], as well as assortative community structure [8] and a densely interconnected core [58].

How do signals traverse neural circuits and what types of neuronal populations do they encounter along the way? The sequence of neurons and neuronal populations that a signal passes through presumably transforms the nature of the signal itself and its downstream effect [1, 4, 24, 41, 42, 53, 64]. For example, signals exchanged between closely clustered and functionally-aligned populations may be relatively unchanged, whereas signals exchanged between anatomically- and functionally-distant populations may be enriched or diversified [5]. A simple way to infer potential signal trajectories in a network is the topological shortest path (hereafter simply referred to as a “path”) [2, 58]. For many classes of networks, including brain networks, decentralized communication mechanisms may also take advantage of shortest paths without any knowledge of the global topology, including diffusion [20] and navigation [54]. Thus, paths connecting pairs of nodes trace out unique motifs along their trajectory, meaning that the nature of communication between any two regions is subject to the underlying structure [4, 24, 40].

Signal propagation is likely to be constrained by the hierarchical organization of cortical circuits. Evidence from classical anatomy and modern neuroimaging points to a continuous sensory-fugal hierarchy, spanning unimodal to transmodal cortex [23, 38]. This continuous axis or gradient can be observed in the functional architecture of the cortex [36], running parallel to gradients in intracortical myelin [30, 48], cortical thickness [61], gene transcription [9, 19], excitation-inhibition ratios [62] and intrinsic temporal time scales [33, 44]. The influence of these multi-modal gradients on signaling and communication in structural networks is a key question in systems neuroscience [53, 60].

Here we investigate how the functional hierarchy shapes the propagation of signals. We reconstruct paths on structural networks and trace their trajectories through the unimodal-transmodal gradient. We find that the hierarchical organization of the cerebral cortex constrains path trajectories, such that most paths follow a canonical bottom-up (ascending the hierarchy) or topdown (descending the hierarchy) trajectory. Importantly, we find that paths may potentially reverse direction in attention networks. Altogether, we find that the hierarchical organization of cortical circuits imposes a communication space on the structural network, potentiating some types of signal propagation patterns while attenuating others.

## RESULTS

The results are organized as follows. We first develop a methodology to trace signal trajectories through the putative unimodal-transmodal hierarchy. We then investigate the extent to which signal flows conform to the hierarchical organization of the cortex, and instances where they diverge. Finally, we consider whether information about hierarchical position is sufficient to sustain a decentralized navigation-like communication process. Data sources include (see *Materials and Methods* for detailed procedures):

- *Structural connectivity*. Structural and functional connectivity were derived from *N* = 66 healthy participants (source: Lausanne University Hospital). Structural connectivity was reconstructed using diffusion spectrum imaging and deterministic streamline tractography. A consistency- and length-based procedure was then used to assemble a group-representative weighted structural connectivity matrix [7, 39, 40].
- *Functional connectivity*. Functional connectivity was estimated in the same individuals using resting-state functional MRI (rs-fMRI). A functional connectivity matrix was constructed using pairwise Pearson correlations among regional time courses. A group-average functional connectivity matrix was then estimated as the mean connectivity of pair-wise connections across individuals.

We trace path motifs between all possible source-target node pairs on the weighted structural network (Fig. 1; for a conceptually similar approach, see [58]). We label nodes according to two different nomenclatures: hierarchical position and intrinsic network affiliation [65]. Hierarchical position is defined as the first principal connectivity gradient of the diffusion map embedding over the FC matrix [36] (see *Materials and Methods*). The continuous embedding vector spans a putative hierarchy, where lower values correspond to unimodal regions and greater values correspond to transmodal regions. We use the empirical cumulative distribution function of the first gradient to bin nodes into ten classes of equal size. We enumerate the classes from 1 to 10, where 1 corresponds to unimodal cortex and 10 to transmodal cortex.

**Figure 1.**
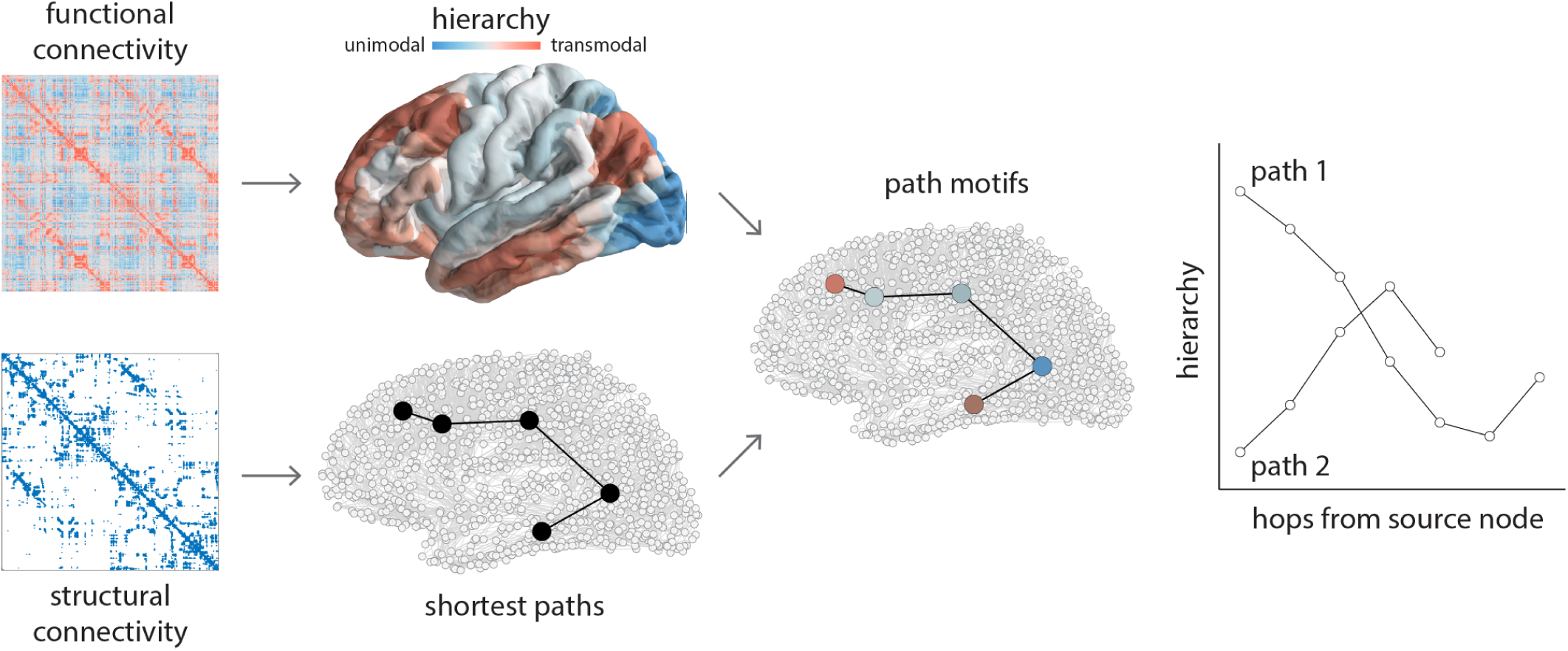
Tracing communication paths through cortical hierarchies. Structural and functional networks are reconstructed from diffusion weighted MRI and resting state functional MRI, respectively. Shortest paths between all pairs of nodes are computed for structural networks using the Floyd-Warshall algorithm [18, 52, 63]. A cortical hierarchy is recovered from functional networks using diffusion map embedding [11]. The first eigenvector is used to label nodes according to their position in the putative unimodal-transmodal hierarchy [36]. Sequences of nodes encountered along a path are labelled by their hierarchical position, tracing out path motifs. Note that some paths are longer and some are shorter, some paths smoothly ascend or descend through the hierarchy, and some paths reverse their trajectory one or more times en route to the target node.

### Path motifs follow hierarchies

We first investigate how path motifs map onto the putative unimodal-transmodal hierarchy. For a given source and target class, we consider all possible paths between the constituent source nodes and target nodes. We then compute the mean hierarchical position of every step encountered along the paths. Fig. 2 shows the mean path motifs originating from a low (class 2), intermediate (class 6) and high (class 9) level source region, to the same three regions as targets. Colours distinguish paths of different lengths.

**Figure 2.**
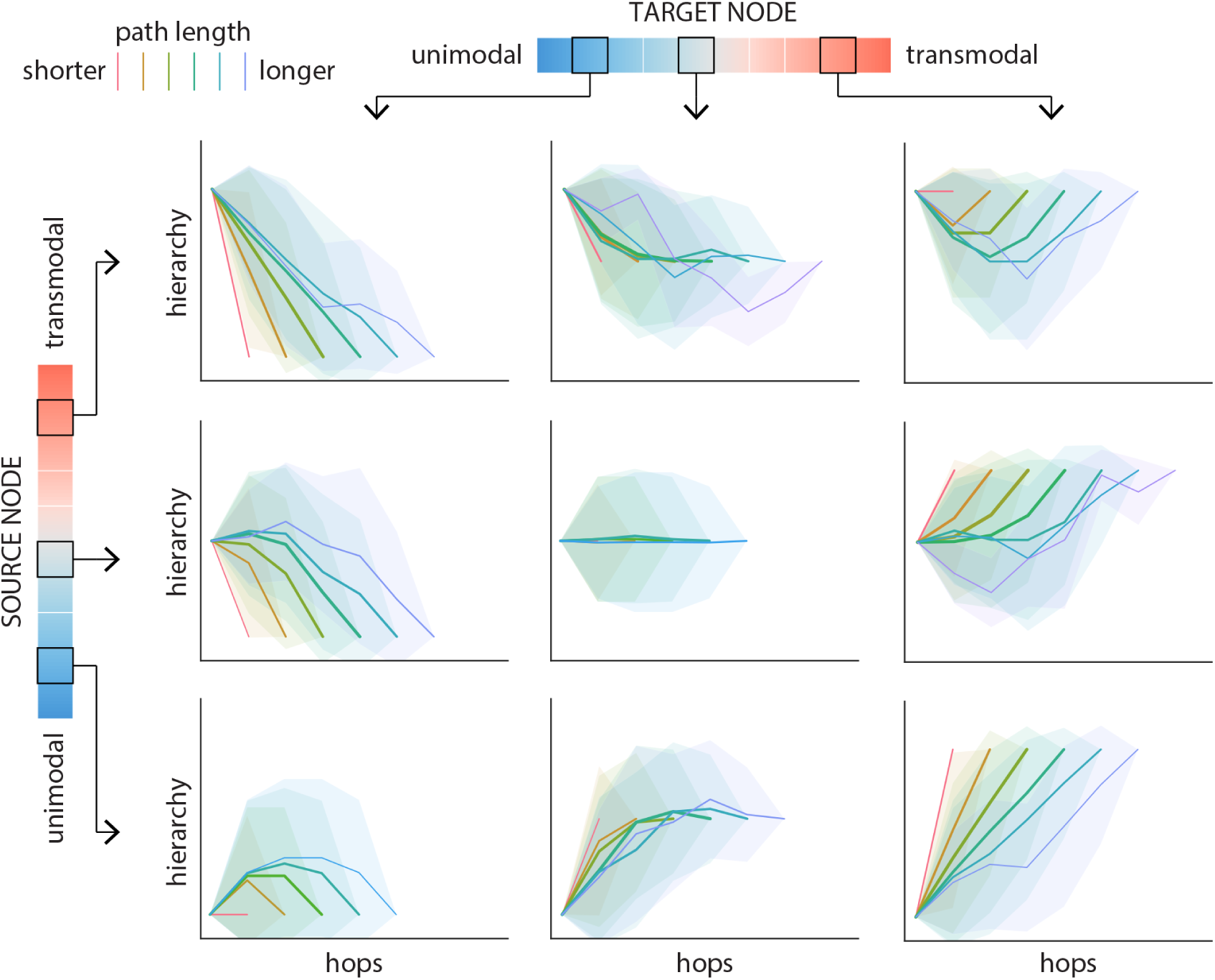
Path motifs. For each source-target pair, nodes along the corresponding path are labeled according to their position on the unimodal-transmodal cortical hierarchy. Hierarchy values are binned into 10 equally-sized levels, where level 1 corresponds to unimodal cortex and level 10 corresponds to transmodal cortex. Paths motifs are shown for three levels of source nodes (2, 6 and 9; rows) and three levels of target nodes (2, 6 and 9; columns). Each plot shows the mean path motifs: path position (hop) is shown on the x-axis and the hierarchical level of the node at each hop is shown on the y-axis. Paths are stratified according to their length, such that warmer colours indicate shorter paths and colder colours indicate longer paths. Shaded regions indicate 95% confidence intervals. Fig. S1 shows the corresponding results for a label-permuting null model.

In general, path motif shape depends on the relative hierarchical position of the source and target nodes (Fig. 2; rows and columns, respectively). For most paths, motifs smoothly transition through the hierarchy, but there exist systematic differences in the nature of the transitions. Paths traversing a larger difference in the hierarchy (e.g. from class 2 to class 9) tend to follow a more monotonic trajectory. Conversely, when the source and the target nodes occupy the same or neighbouring positions in the hierarchy, paths are more likely to follow a U-shape, effectively taking detours to intermediate parts of the hierarchy (e.g. when source and target is class 2). These trajectories are in contrast to trajectories recovered from null networks with randomized hierarchy labels, wherein paths traverse the hierarchy more slowly and less smoothly (Fig. S1).

### Inflection points in communication flow

In the previous section we observed that anatomical connectivity fundamentally shapes how the unimodal-transmodal hierarchy is traversed, promoting some transitions while attenuating others. To investigate how these “pushing” and “pulling” forces shape the communication landscape, we next consider path trajectories at the nodal level. Specifically, we study how the orientation of the flows changes along the course of the journey towards the target, which we quantify by the slope of paths through the hierarchy.

For a given node, we calculate the mean slope of all paths as they pass through that node (Fig. 3a). If, on average, the slope of the paths when passing through a node is positive, this suggests that the transmission is ascending through the hierarchy, from unimodal towards transmodal cortex. Conversely, nodes with negative slopes direct information flow towards areas lower in the cortical hierarchy. Note that, in general, a node could participate in both ascending and descending paths, and this dependent measure reflects the mean flow of information through that node.

**Figure 3.**
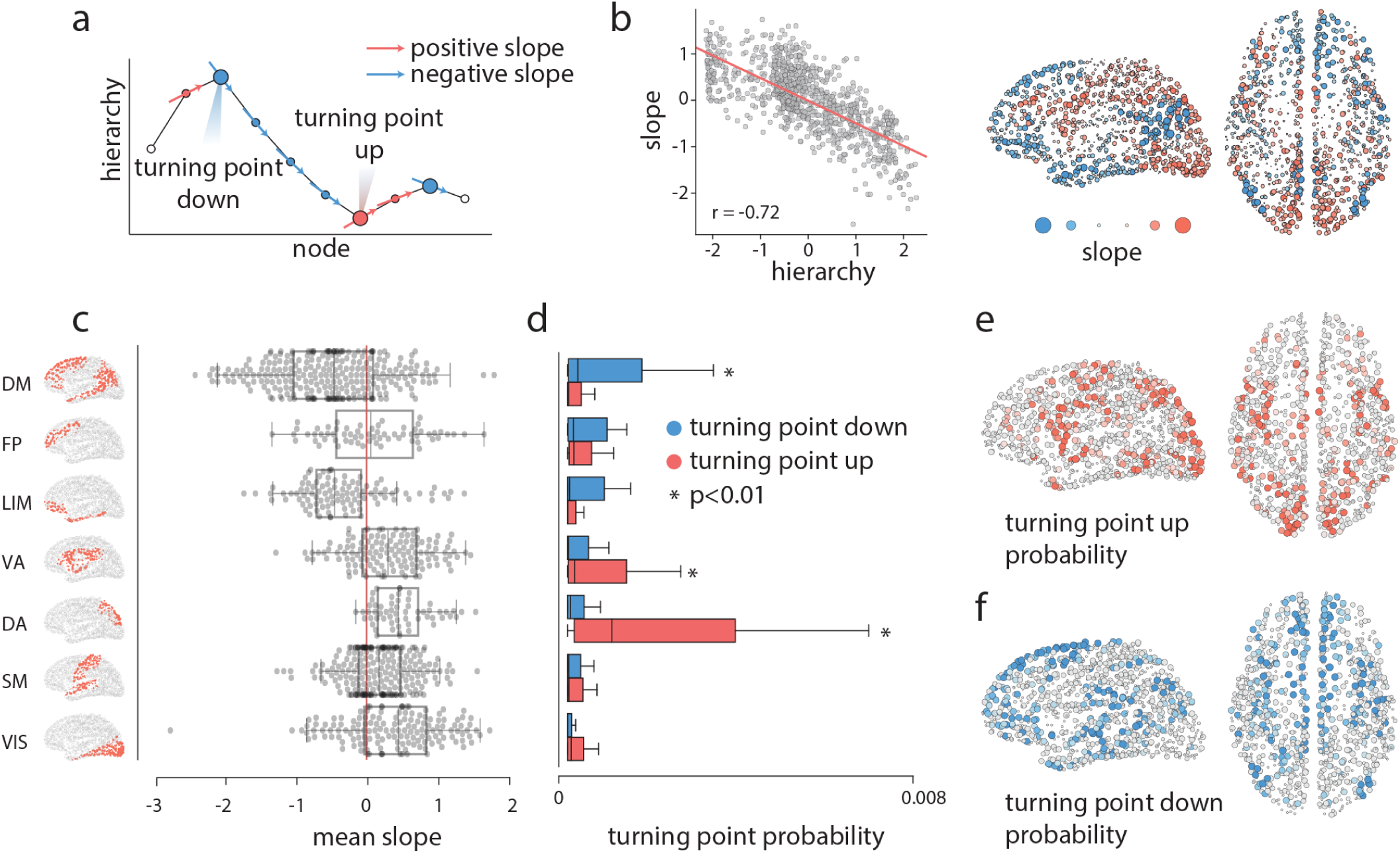
Inflection points in communication flow. As each path traverses the hierarchy, we can infer how individual brain regions direct communication flow. (a) Schematic showing a path motif, where the position along the x-axis indicates a node and the y-axis indicates the hierarchical position of the node. The slope of the curve at each point tells whether the path is ascending or descending in the hierarchy. We denote nodes where slope changes sign as turning points. Nodes where the slope changes from positive to negative are turning points down, and nodes where the slope changes from negative to positive are turning points up. (b) The mean slope of each node (y-axis) is anticorrelated with its position in the hierarchy. The mean slope of each node is shown for every brain region; warm colours indicate positive slopes, cold colours indicate negative slopes. (c) The mean slope for seven intrinsic networks [65]. (d) Mean probability of turning points up and down in seven intrinsic networks. Asterisks indicate values that are statistically significant according a label-permuting null distribution (*p* < 0.01, FDR-corrected). (e) Turning point up probability for individual regions. (f) Turning point down probability for individual regions. Network assignments: DM = default mode, FP = fronto-parietal, LIM = limbic, VA = ventral attention, DA = dorsal attention, SM = somatomotor, VIS = visual.

We first note that the mean slope for a given node is negatively correlated with the node’s position in the unimodal-transmodal hierarchy (Fig. 3b; *r* = −0.72). In other words, nodes that occupy higher positions in the hierarchy tend to direct signal traffic towards nodes lower in the hierarchy, and vice versa. This is consistent with the intuition developed in the previous subsection. Fig. 3b shows that areas that exhibit mainly positive mean slope (i.e. direct information to ascend the hierarchy) are the supplementary motor area, somatomotor cortex and visual cortex. Areas with negative slope (i.e. directing information to descend the hierarchy) are the prefrontal cortex, the posterior parietal cortex, auditory cortex and the inferotemporal cortex. Stratifying these nodes by their membership in intrinsic networks [65], we find mean positive slopes for the visual, somatomotor, dorsal attention, ventral attention and frontoparietal networks, and mean negative slopes for the limbic and default mode networks (Fig. 3c). Given that mean slope is anticorrelated with hierarchical position, for completeness we also regressed out hierarchical position to reveal nodes where slopes (i.e. tendency to direct information) were either greater or lower than expected given their hierarchical position. Fig. S2 shows that these hierarchy-corrected slopes are generally similar, with greater emphasis on dorosolateral prefrontal cortex and medial parietal cortex as regions that direct communication higher in the hierarchy.

The slope of a path traversing a node also allows us to identify areas that re-direct flow direction and promote detours. As we follow a path trajectory, we look for local extremum nodes that reside between slopes with different signs, and tag them as turning points. Depending on the type of extremum, we name the turning points as “turning up” points (local minima) or “turning down” points (local maxima) (Fig. 3a). For example, a turning down point is a node that occupies a relatively higher position in the hierarchy and connects two lower-level brain areas. We first stratify nodes by their membership in intrinsic networks and compute the mean turning point probability for each network. Fig. 3d shows that networks with the greatest probability of turning up paths (i.e. re-directing them to ascend the hierarchy) are the dorsal attention and ventral attention, whereas the default mode network has the greatest turning down probability. At the regional level, regions with the greatest probability of turning up the paths tend to be in attention-related networks, including the supplementary motor area and posterior parietal cortex (Fig. 3e). Conversely, superior and dorsolateral prefrontal, inferotemporal and lateral temporal cortex are the most probable turning down points (Fig. 3f).

### Temporal evolution of communication flow

How does the hierarchical organization of the brain shape communication across time? Given that most communication paths conform to the hierarchical organization of the network, we next ask whether the hierarchy imparts memory on communication processes by exerting influence on the path trajectories. To address this question, we consider the temporal evolution of communication patterns, envisioning the sequence of nodes traversed along a path as a time series.

We explore how the position of a walker traversing the path depends on the positions it occupied previously in its trajectory. Specifically, we compute the probability of going from a node that belongs to the hierarchy level *i* to a node that belongs to hierarchy level *j* in one step as a function of the position over the path (1-hop transitions; Fig. 4a, left). To quantify whether the hierarchical position of a walker depends on its previous hierarchical positions (multi-hop transitions), we measure the transition probability of occupying hierarchy level *i* at step *t*, given that the walker occupied hierarchy level *j* at step *t* − *k*, changing *k* from 1 to the length of the path (multihop transitions; Fig. 4a, right).

**Figure 4.**
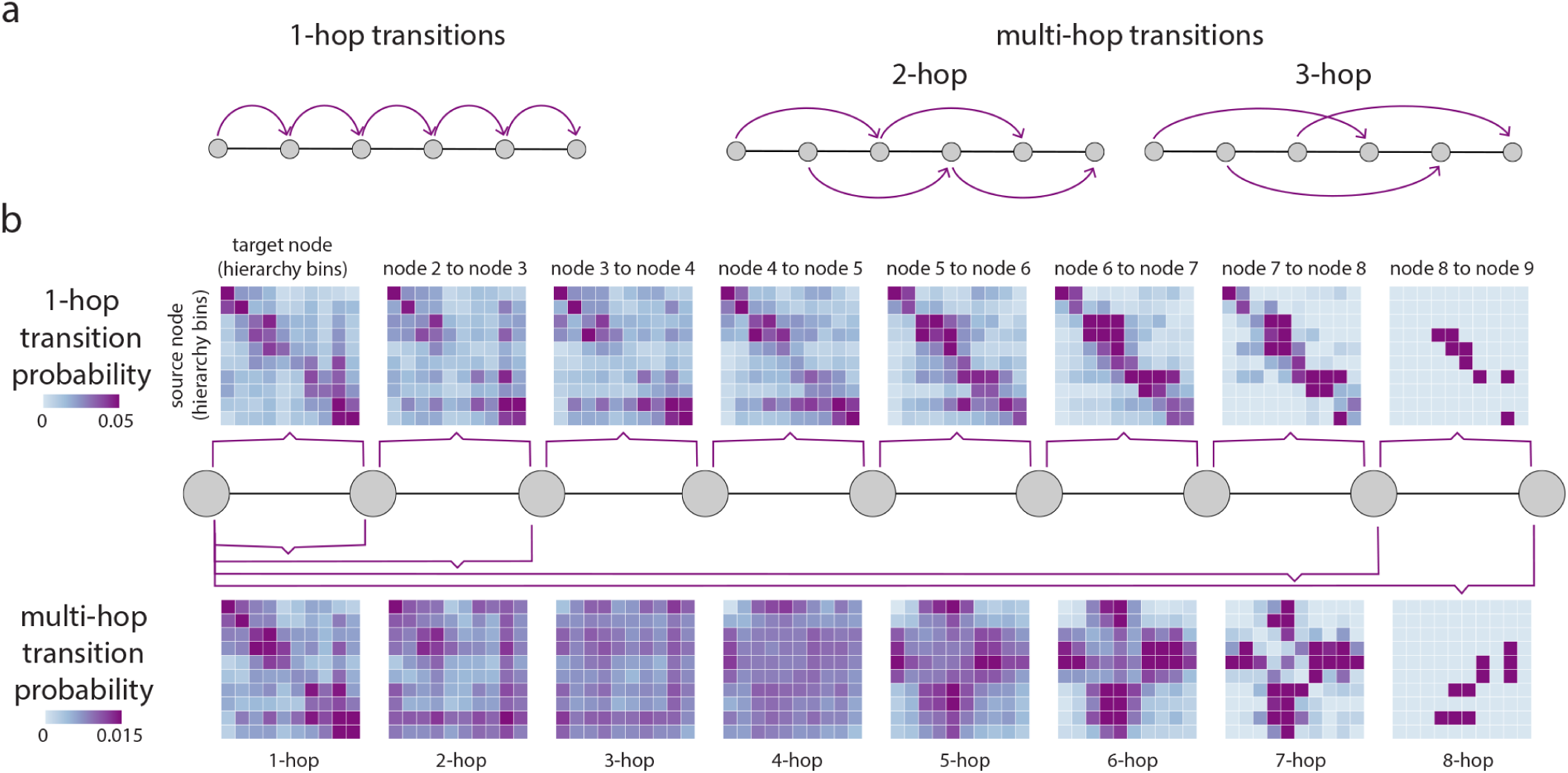
Transition probabilities in communication. As paths traverse the hierarchy, we quantify the probability that the current position of a node in the unimodal-transmodal hierarchy depends on previous positions in the path. (a) With 1-hop transitions we quantify the transition probability of a path going from hierarchy level *i* to level *j* in one step or hop. With multi-hop transitions we quantify the probability of a path going from hierarchy level *i* to level *j* in *k* steps or hops. Thus, in a single path we consider multiple transitions. (b) Nodes are stratified by their hierarchical position in 10 equally-sized bins. Transition probability matrices are shown where source nodes (hierarchy bins) are in the rows, and target nodes (hierarchy bins) are in the columns. The top row shows 1-hop transition probabilities and the bottom row shows multi-hop transition probabilities. Transitions are shown for paths of up to length 9, corresponding to the diameter of the network. Note that the values in the matrices display mean probabilities over multiple paths, hence the rows do not necessarily sum to 1 (see *Materials and Methods* for more detail).

Fig. 4b (top) shows 1-hop transition probabilities, as in a first-order Markov process. Nodes are stratified according to their position in the hierarchy, with source nodes in the rows and target nodes in the columns. Mean transition probabilities are greatest over the diagonal, favouring the transmission of a walker to neighbouring positions in the hierarchy, repeating the theme from the previous two subsections. Fig. 4 (bottom) shows average probabilities for transitions (memories) over 2 steps or more. For memories 2, 3 and 4 steps away, transitions become more uniform, meaning that the probability of occupying current position *j* does not depend greatly on the position it occupied 2, 3 or 4 steps before. For greater memory values of 5, 6 and 7, there is an emergence of transitions between lower and intermediate, and higher and intermediate hierarchy levels, with almost zero probability of a transition between lower and higher levels.

Altogether, we find that most 1-hop transitions smoothly follow the hierarchy, but that communication over longer trajectories is biased towards some levels of the hierarchy and away from others, particularly if the starting point is at intermediate levels. In other words, the nodes visited by a walker earlier in the trajectory may exert influence on transition probabilities later in the trajectory.

### Navigation via hierarchies

Given that communication paths closely align with the hierarchy of the network, we finally ask whether it is possible to recapitulate the path architecture of the network by following the hierarchy. We focus on navigation, a decentralized communication mechanism in which a signal is forwarded to the connected neighbour that is closest in some distance to the target. This distance is defined with respect to an underlying metric space, with the simplest such space being the three-dimensional space that nodes are physically embedded in. For example, previous work has demonstrated that it is possible to recapitulate the shortest path architecture by forwarding signals to nodes that are physically closest to the target node [54]. Decentralized mechanisms such as navigation are intuitively appealing as they do not assume that signals or nodes possess knowledge of the global topology [3, 54].

We therefore investigate whether signals could recapitulate the path structure of the network if they are forwarded to the neighbour closest to the target node in the unimodal-transmodal hierarchy. To quantify navigation as a communication process we measure the proportion of paths that are successfully recovered (success ratio; *S*_*R*_). Given the importance of spatial embedding, we identify regions for which navigation success improves when hierarchy information is taken into account rather than only spatial factors. To operationalize a node’s proximity to the target we use a linear combination of Euclidean distance in three-dimensional physical space and distance in “hierarchy space” weighted by the parameter *β* as:

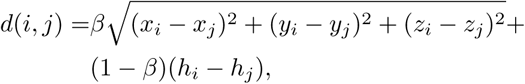

where *d*(*i, j*) is the combined distance between nodes *i* and *j*, (*x*_*i*_, *y*_*i*_, *z*_*i*_) gives the position of node *i* in three-dimensional Euclidean space and *h*_*i*_ gives the position of node *i* in one-dimensional hierarchical space. For each pair of nodes we measure the success ratio as a function of *β*, tuning *β* from 0 to 1, and find the *β* that maximizes the navigation success. When *β* is valued close to 1, paths originating from the node are better recovered using spatial proximity compared to hierarchical proximity; the opposite is true when *β* is valued close to 0 (Fig. 5a; see *Materials and Methods* for more detail).

**Figure 5.**
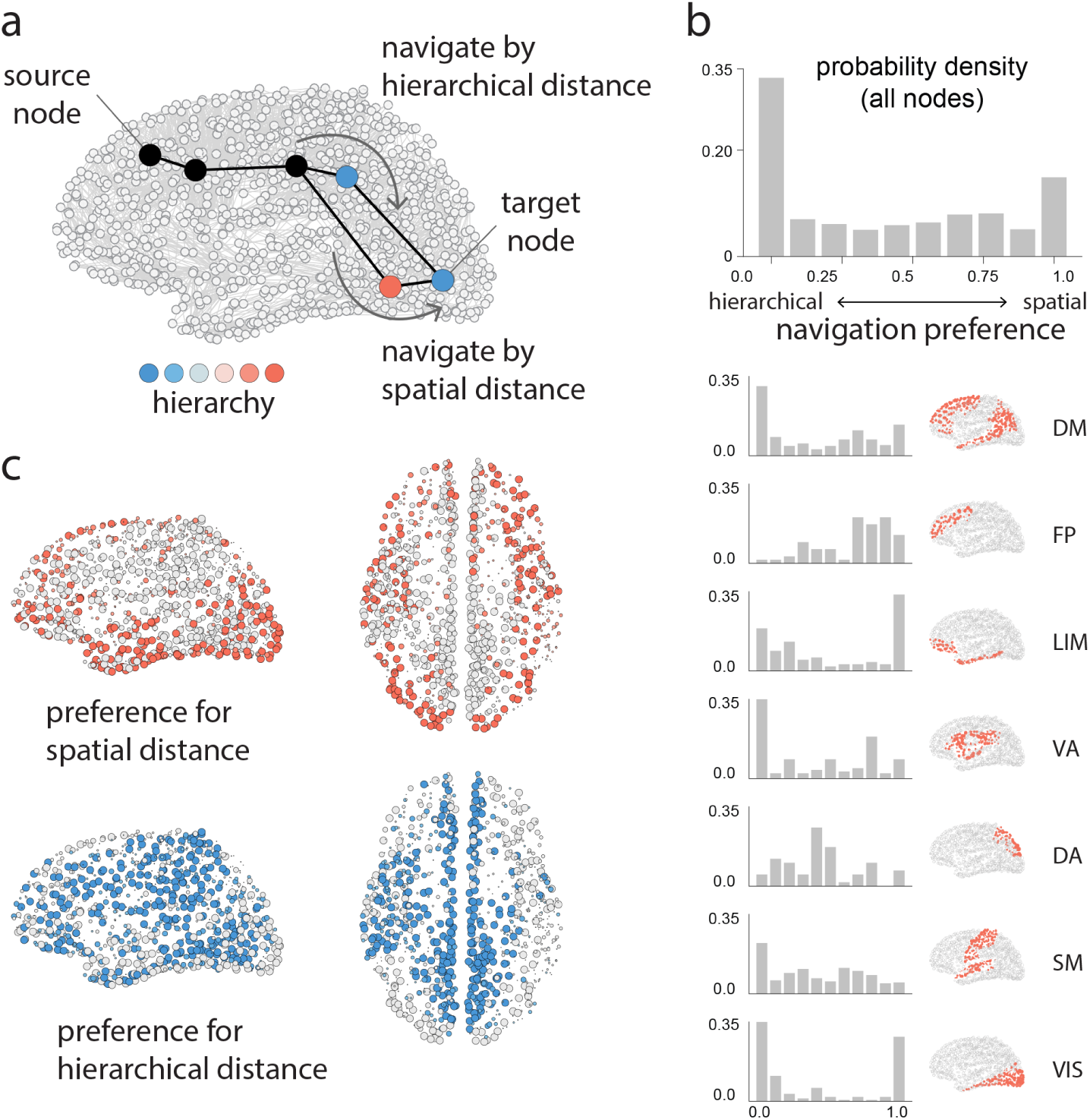
Navigation via hierarchical proximity. We evaluate the extent to which shortest paths can be recapitulated by an agent who is aware of the three-dimensional spatial positions of the nodes (spatial navigation) and/or the hierarchical positions of the nodes (hierarchical navigation), but not the topology of the network. (a) Schematic showing a putative path from source target, where an agent occupies the third node in the path. Nodes are coloured according to their position in the hierarchy. If the agent navigates using spatial information, it will transition to the neighbour that is physically closest to the target (bottom). If the agent navigates using hierarchical information, it will transition to the neighbour that is hierarchically closest to the target (top). We derive the navigation preference of each node (*β* parameter) as the source of information that maximizes recapitulation of shortest paths [45, 46, 54]. When *β* is valued close to 1, paths originating from the node are better recovered using spatial proximity compared to hierarchical proximity; the opposite is true when *β* is valued close 0. (b) Histogram of *β* values across all nodes in the network. Histograms are also shown for seven intrinsic networks. (c) Individual brain regions are coloured by their preference for spatial navigation (warm colours; *β* > 0.8) or hierarchical navigation (cold colours; *β* < 0.2). Network assignments: DM = default mode, FP = fronto-parietal, LIM = limbic, VA = ventral attention, DA = dorsal attention, SM = somatomotor, VIS = visual.

Fig. 5b shows the distribution of the *β* parameters that maximize navigation success for each source-target pair. The distribution is bimodal, with prominent peaks at the extremes, suggesting that most nodes have a strong preference for either hierarchical or spatial navigation. Stratifying nodes by membership in intrinsic networks, we find that each network exhibits a unique fingerprint, with some showing a preference for spatial navigation (frontoparietal), others for hierarchical navigation (default), and others a mix between the two (dorsal attention). Fig. 5c shows parts of the visual system, lateral temporal cortex and dorsolateral prefrontal cortex exhibit a strong preference for spatial navigation (red; *β* > 0.8), while medial parietal cortex, medial prefrontal cortex and left temporo-parietal cortex exhibit a strong preference for hierarchical navigation (blue; *β* < 0.2).

### Relation with simple measures

In the present report we derived four dependent measures based on the concept of path motifs (slope, tuning point up/down probability and navigation preference). For completeness, we assess the extent to which these node-level can be related to simpler measures. Fig. S3 shows linear regressions comparing each of the four path motif measures (rows) with simpler network measures computed from structural and functional connectivity matrices (columns). From the structural network we compute betweness, closeness, clustering, degree and mean edge length. From the functional network we compute strength and participation coefficient (relative to the intrinsic network partition reported by Yeo and colleagues [65]). As expected, we find weak to moderate correlations with path-based measures (betweenness, closeness) and with degree, consistent with the notion that most centrality measures are correlated with each other [47]. In addition, we find a positive correlation between participation and mean slope (*r* = 0.52), suggesting that nodes with more diverse connection profiles are more likely to direct communication towards the apex of the unimodal-transmodal hierarchy. In sum, we find that the four path motif-derived measures are correlated with some simpler measures, but cannot be wholly predicted from any one such measure.

## DISCUSSION

In the present report we asked how signals travel on brain networks and what types of nodes they potentially visit en route. We traced individual path motifs to investigate the propensity of communication paths to explore the putative unimodal-transmodal cortical hierarchy. We find that the architecture of the network promotes signaling via the hierarchy, suggesting a link between the structure and function of the network. Importantly, we also find instances where detours are promoted, particularly as paths traverse attention-related networks. Finally, information about hierarchical position aids navigation in some parts of the network, over and above spatial location. Altogether, the present results touch on a number of emerging themes in network neuroscience, including the nature of structure-function relationships, network communication and the role of cortical hierarchies.

The most prominent observation is that most paths closely follow the cortical hierarchy, traveling smoothly from unimodal to transmodal cortex and vice versa (Figs. 2). This is reminiscent of the notion that much of signal traffic follows a sequential bottom-up or top-down trajectory, potentiating direct stimulus-response patterns [64]. At the same time, the architecture of the network occasionally serves to re-direct signal traffic and promote detours. In particular, the default network appears to be the most likely mediator between areas lower in the cortical hierarchy, while the dorsal attention and ventral attention networks act as mediators between areas higher in the hierarchy (Fig. 3). The finding that attention networks are anatomically positioned to re-direct signal traffic resonates with moderns theories of how attention and control networks promote fluid transitions between segregated and integrated states, promoting adaptive reconfiguration during rest and task [13, 17, 43, 55].

These results suggest that the architecture of the network promotes behaviourally-relevant communication patterns, and that the functional properties of individual areas are fundamentally related to their anatomical network embedding. Indeed, multiple studies point to the idea that the topology of brain networks endows individual regions with specific functional attributes. For example, functional properties depend on local connectional profiles, including motif composition [21], asymmetry [34], length [60] and weight distribution [37]. At the global level, anatomical segregation, most prominently observed in the unimodal visual and somatosensory cortices, promotes specialized processing. Conversely, polysensory association cortex is anatomically better integrated in the connectome, potentially allowing information to be sampled from multiple parts of the network [40, 41, 58]. A recent report demonstrated that regions at the top of the hierarchy are better positioned to act as “receivers” of information, while regions at the bottom are better positioned to act as “senders” [53]. The present results build on this literature, showing that the default and attentional networks present links between parts of the cortical hierarchy.

In shaping communication patterns, network architecture may also impart memory on signal traffic, such that transitions depend not only on the current position of the signal, but also on positions they previously occupied in the hierarchy in their journey (Fig. 4). We find that most transitions between nodes that occupy neighbouring positions in the hierarchy are memoryless, but that transitions across disparate levels are not. The phenomenon of network structure imposing non-Markovian network flows is also observed in other complex systems, such as air passenger flows and journal citation flows [51]. In the brain, the functional consequences of this phenomenon may be the well-studied hierarchical organization of time scales and temporal receptive windows. Numerous studies point to the idea that information accumulates over time across the cortical hierarchy, such that the temporal dynamics of higher-order regions unfold over slower time scales [33, 44], manifesting as a preference for long-range contextual information [29]. Understanding the precise link between network structure and the hierarchy of intrinsic time scales remains a major question for future research [15, 22].

More generally, these results open new questions about how broad spatial gradients in synaptic connectivity, cytoarchitecture and molecular composition interact with macroscale network topology [19, 36, 60, 62]. Current graph models of brain networks assume that all nodes are the same, but as signals propagate through the network, they pass through a series of heterogeneous neural circuits and populations [1, 56]. Each stage in the trajectory may entail transformations that modern methods in network neuroscience do not consider. For example, the majority of path-based metrics consider the total lengths of paths between areas, but not the identity of nodes traversed during the path. By drawing path motifs through maps annotated by molecular and cellular data, the present methodology permits closer investigation into how local attributes of nodes may influence communication in the network.

Note that the present method only traces out the shortest paths in the network, but this does not preclude the possibility that communication takes place via mechanisms that are unaware of the shortest paths in the network [2, 3, 56]. Several recent reports point to diffusion-like and navigation-like processes as potentially more biologically-realistic alternatives [20, 26, 42, 54], as they do not assume that signals possess knowledge of the global topology. At the same time, multiple studies suggest that shortest paths in brain networks are readily accessible by both diffusion [20] and spatial navigation (greedy routing) [54], without knowledge of the global topology. Thus, shortest paths are an important substrate for communication, even if communication does not occur via shortest path routing per se [3]. We envision that future studies will consider diffusion and navigation trajectories, analogous to the approach we took with shortest paths.

We close by noting important methodological considerations. Although the present networks are derived from a consensus of 66 participants with high-quality imaging [7], there are several limitations. First, structural networks were reconstructed using diffusion weighted MRI and computational tractography, a technique that results in systematic false positives and false negatives [14, 35, 57]. Second, the technique cannot be used to resolve the direction of white matter projections, which means that some paths recovered from the network may not exist. Third, the present reconstruction only includes cortical regions, leaving out important topological contributions from the subcortex and cerebellum. Network communication is undoubtedly shaped by both sets of structures, and future studies should consider subcortical-cortical and cerebellar-cortical signal traffic.

In summary, we develop a simple framework to trace communication patterns in brain networks. We show that the putative unimodal-transmodal hierarchy shapes the propagation of signals, imparting behaviourally-relevant communication patterns. The present work highlights the importance of considering sequences of nodes encountered during signaling, and the role they might play in network-wide communication.

## METHODS

### Data acquisition

A total of N = 66 healthy young adults (16 females, 25.3 ± 4.9 years old) were scanned at the Department of Radiology, University Hospital Center and University of Lausanne. The scans were performed in 3-Tesla MRI scanner (Trio, Siemens Medical, Germany) using a 32-channel head-coil. The protocol included (1) a magnetization-prepared rapid acquisition gradient echo (MPRAGE) sequence sensitive to white/gray matter contrast (1 mm in-plane resolution, 1.2 mm slice thickness), (2) a diffusion spectrum imaging (DSI) sequence (128 diffusion-weighted volumes and a single b0 volume, maximum b-value 8000 s*/*mm_2_, 2.2 × 2.2 × 3.0 mm voxel size), and (3) a gradient echo EPI sequence sensitive to BOLD contrast (3.3 mm in-plane resolution and slice thickness with a 0.3 mm gap, TR 1920 ms, resulting in 280 images per participant). Participants were not subject to any overt task demands during the fMRI scan.

### Structural network reconstruction

Grey matter was parcellated into 68 cortical nodes according to the Desikan-Killiany atlas [16]. These regions of interest were then further divided into 1000 approximately equally-sized nodes [10]. Structural connectivity was estimated for individual participants using deterministic streamline tractography. The procedure was implemented in the Connectome Mapping Toolkit [12], initiating 32 streamline propagations per diffusion direction for each white matter voxel. Structural connectivity between pairs of regions was defined as the number of streamlines normalized by the mean length of streamlines and mean surface area of the two regions, termed fiber density [27]. This normalization compensates for the bias toward longer fibers during streamline reconstruction, as well as differences in region size.

To mitigate concerns about inconsistencies in reconstruction of individual participant connectomes [31, 57], as well as the sensitive dependence of network measures on false positives and false negatives [66], we adopted a group-consensus approach [7, 14, 50]. In constructing a consensus adjacency matrix, we sought to preserve (a) the density and (b) the edge length distribution of the individual participants matrices [6, 7, 40]. We first collated the extant edges in the individual participant matrices and binned them according to length. The number of bins was determined heuristically, as the square root of the mean binary density across participants. The most frequently occurring edges were then selected for each bin. If the mean number of edges across participants in a particular bin is equal to *k*, we selected the *k* edges of that length that occur most frequently across participants. To ensure that inter-hemispheric edges are not under-represented, we carried out this procedure separately for inter- and intra-hemispheric edges. The binary density for the final whole-brain matrix was 2.17%. The weight associated with each edge was then computed as the mean weight across all participants.

### Functional network reconstruction

Functional MRI data were pre-processed using procedures designed to facilitate subsequent network exploration [49]. FMRI volumes were corrected for physiological variables, including regression of white matter, cerebrospinal fluid, as well as motion (three translations and three rotations, estimated by rigid body co-registration). BOLD time series were then subjected to a lowpass filter (temporal Gaussian filter with full width half maximum equal to 1.92 s). The first four time points were excluded from subsequent analysis to allow the time series to stabilize. Motion “scrubbing” was performed as described by Power and colleagues [49]. The data were parcellated according to the same atlas used for structural networks [10]. Individual functional connectivity matrices were defined as zero-lag Pearson correlation among the fMRI BOLD time series. A group-consensus functional connectivity matrix was estimated as the mean connectivity of pair-wise connections across individuals.

### Diffusion map embedding

Diffusion map embedding is a nonlinear dimensionality reduction algorithm [11]. The algorithm seeks to project a set of embeddings into a lower-dimensional Euclidean space. Briefly, the similarity matrix among a set of points (in our case, the correlation matrix representing functional connectivity) is treated as a graph, and the goal of the procedure is to identify points that are proximal to one another on the graph. In other words, two points are close together if there are many relatively short paths connecting them. A diffusion operator, representing an ergodic Markov chain on the network, is formed by taking the normalized graph Laplacian of the matrix. The new coordinate space is described by the eigenvectors of the diffusion operator. We set the diffusion rate *α* = 1 and the variance of the Gaussian used in affinity computation *σ* = 1. The procedure was implemented using the Dimensionality Reduction Toolbox (https://lvdmaaten.github.io/drtoolbox/) [59].

### Shortest path retrieval

Structural connectivity was encoded as an undirected weighted graph *G* ≡ {*V, W*} comprised of nodes *V* = {*v*_1_, *v*_2_, …*v*_*n*_} and a matrix of fiber density values *W* = [*w*_*ij*_], valued on the interval [0,1]. To recover shortest paths, we first define a topological distance measure. The weighted adjacency matrix was transformed from a connection weights to connection lengths matrix using the transform *L* = −*log*(*W*), such that connections with greater weights are mapped to shorter lengths [20]. Note that other transformations are also possible, including *L* = 1*/W*. The drawback of this transform is that it generates highly skewed distributions of lengths *L*. As a result, a small number of connections are valued much more than the rest, and they are disproportionately represented in shortest paths. The logarithmic transform controls for this, yielding log-normal distributions of *L* [2]. Weighted shortest paths were recovered using the Floyd-Warshall algorithm [18, 52, 63] (*Brainconn* Python Toolbox). Note that in many types of networks there may exist multiple shortest paths between two nodes (edge-disjoint or not); in our network this was not the case as we computed weighted shortest paths, yielding unique paths between all source-target pairs.

### Inflection points

A shortest path is defined as a sequence of nodes *v*_1_, *v*_2_, …, *v*_*n*_ where *v*_*i*_ is the node that was at the i-th step of a path of length *n*. Each node has an associated hierarchical position (value), so we assign the difference between the hierarchy values *h*_*v*_*i, h*_*v*_*i*+1 as the slope of the path through node *v*_*i*_. Once all slopes are assigned, we compute the mean over all paths for each node.

### Network navigation

We measured navigation by simulating an agent or walker that traverses the network from source node *i* to target node *j*. The agent has no knowledge of the global topology; instead, they hop towards neighbours who are closest to the target node in some underlying metric space. Across all source-target pairs, we measure the proportion of paths that are successfully recovered (success ratio; *S*_*R*_) [54]. To operationalize a node’s proximity to the target we use a linear combination of the Euclidean distances in three-dimensional physical space and in hierarchy space weighted by the parameter *β* as:

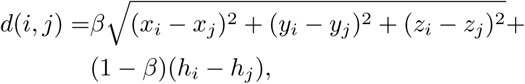

where *d*_*i,j*_ is the combined distance between nodes *i* and *j*, (*x*_*i*_, *y*_*i*_, *z*_*i*_) is a node’s position in Euclidean space and *h*_*i*_ is a node’s position in hierarchy space. Both values were normalized to lie in the interval [0,1]. Given that navigation is a deterministic model, we calculated, for each node as a source, the success ratio as a function of *β*. The resulting curves showed a global trend preferring Euclidean distance, with optimal *β* values close to 0.8, consistent with previous reports [54]. In each node, however, there exists substantial variance across *β* values, showing a changing preference deviating from global trend. To better capture this preferences for individual nodes, we detrend the mean success ratio and select for each source node the *β* that maximizes the detrended success ratio (Fig. S4).

### Transition probabilities

We define the one-hop transition probability matrix **T** as a function of the position *t*, where *t* = 1, …, *Diam*(*G*_*SC*_) − 1. For each position there was one matrix **T**(*t*) defined as

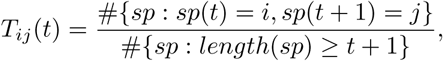

where *i* and *j* represent the hierarchy bins, thus (*i, j*) ∈ {*h*_1_, …, *h*_10_}. The expression {# *sp*: ○}represents the number of shortest paths, from the set of all shortest paths, that satisfy the condition ○.

For multi-hop transition probabilities, we define the transition matrix between hierarchy bins as a function of hops *k*, denoted as **M**(*k*), where *k* = 1, …, *Diam*(*G*_*SC*_) − 1. For each hop length *k* the matrix **M**(*k*) was defined as

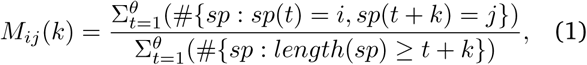

where *i, j* ∈ {*h*_1_, …, *h*_10_}, *θ* = *length*(*sp*) − *k, θ* ∈ [1, *Diam*(*G*_*SC*_) − 1]

### Null model

The critical question underlying all reported analyses is the link between structural connectivity and hierarchical position. To assess this question, we used a null model that randomly permutes the hierarchical label of nodes while preserving their topological and spatial embedding (2,000 repetitions) [67], embodying the null hypothesis that there is no relationship between structural topology and functional hierarchy labels.

## ACKNOWLEDGMENTS

The authors thank Dr. Alessandra Griffa for collecting, preprocessing and sharing the neuroimaging dataset. We acknowledge the Department of Psychiatry of Lausanne University Hospital and particularly Professor Philippe Conus, Professor Kim Do Cuenod, Raoul Jenni and Martine Cleusix for having helped with the recruitment process of the study volunteers. The authors thank Ross Markello, Golia Shafiei, Vincent Bazinet, Laura Suarez and Justine Hansen for helpful comments on the manuscript. This research was undertaken thanks in part to funding from the Canada First Research Excellence Fund, awarded to McGill University for the Healthy Brains for Healthy Lives initiative. BM acknowledges support from the Natural Sciences and Engineering Research Council of Canada (NSERC Discovery Grant RGPIN #017-04265), from the Canada Research Chairs Program and from the Fonds de recherche du Québec - Santé (Chercheur Boursier).

## Data availability

The processed dataset (structural and functional matrices) is available at DOI: 10.5281/zenodo.2872624 [25].

---

**Figure S1.**
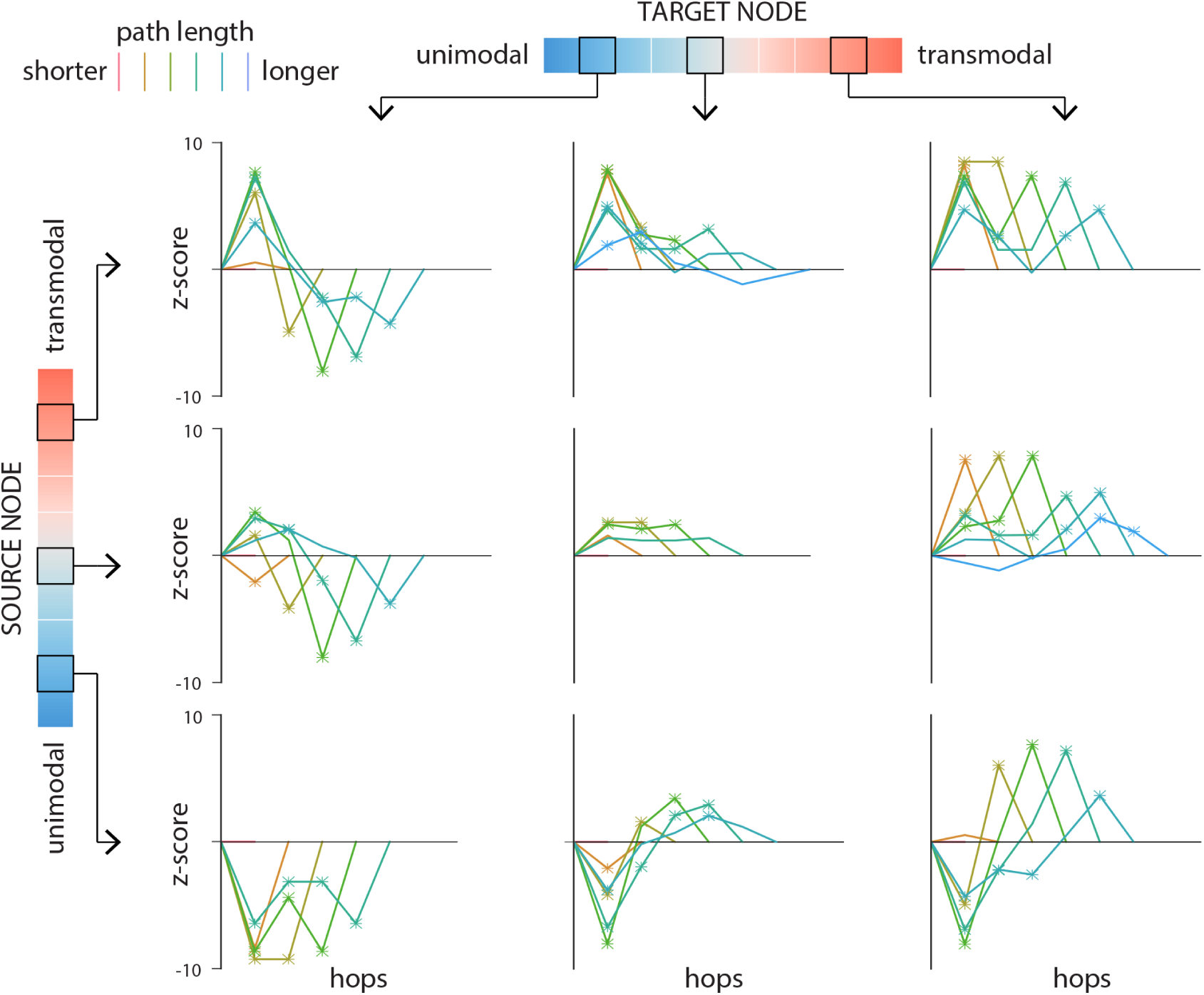
Path motifs null model. For each source-target pair, nodes along the corresponding path are labeled according to their position on the unimodal-transmodal cortical hierarchy. Hierarchy values are binned into 10 equally-sized levels, where level 1 corresponds to unimodal cortex and level 10 corresponds to transmodal cortex. Paths motifs are shown for three levels of source nodes (2, 6 and 9; rows) and three levels of target nodes (2, 6 and 9; columns). Each plot shows the z-score of the mean path motif relative to a label-permuting null mode. Path position (hop) is shown on the x-axis. Paths are stratified according to their length, such that warmer colours indicate shorter paths and colder colours indicate longer paths. Points denoted by asterisks indicate *p* < 0.05.

**Figure S2.**
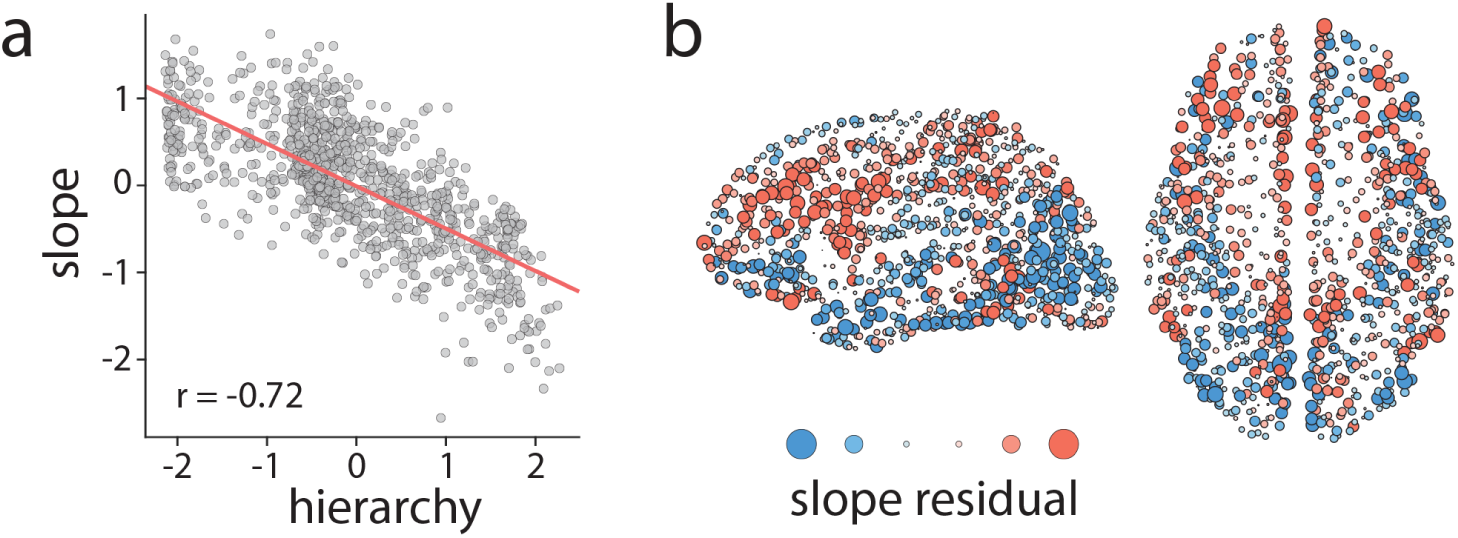
Mean slope before and after correcting for hierarchical position. (a) The mean path slope shown in Fig. 3 is anticorrelated with hierarchical position. (b) Mean slope after linearly regressing out hierarchical position. Warm colours indicate regions where paths ascend the hierarchy more than expected given their hierarchical position; cold colours indicate regions where paths descend the hierarchy more than expected given their hierarchical position.

**Figure S3.**
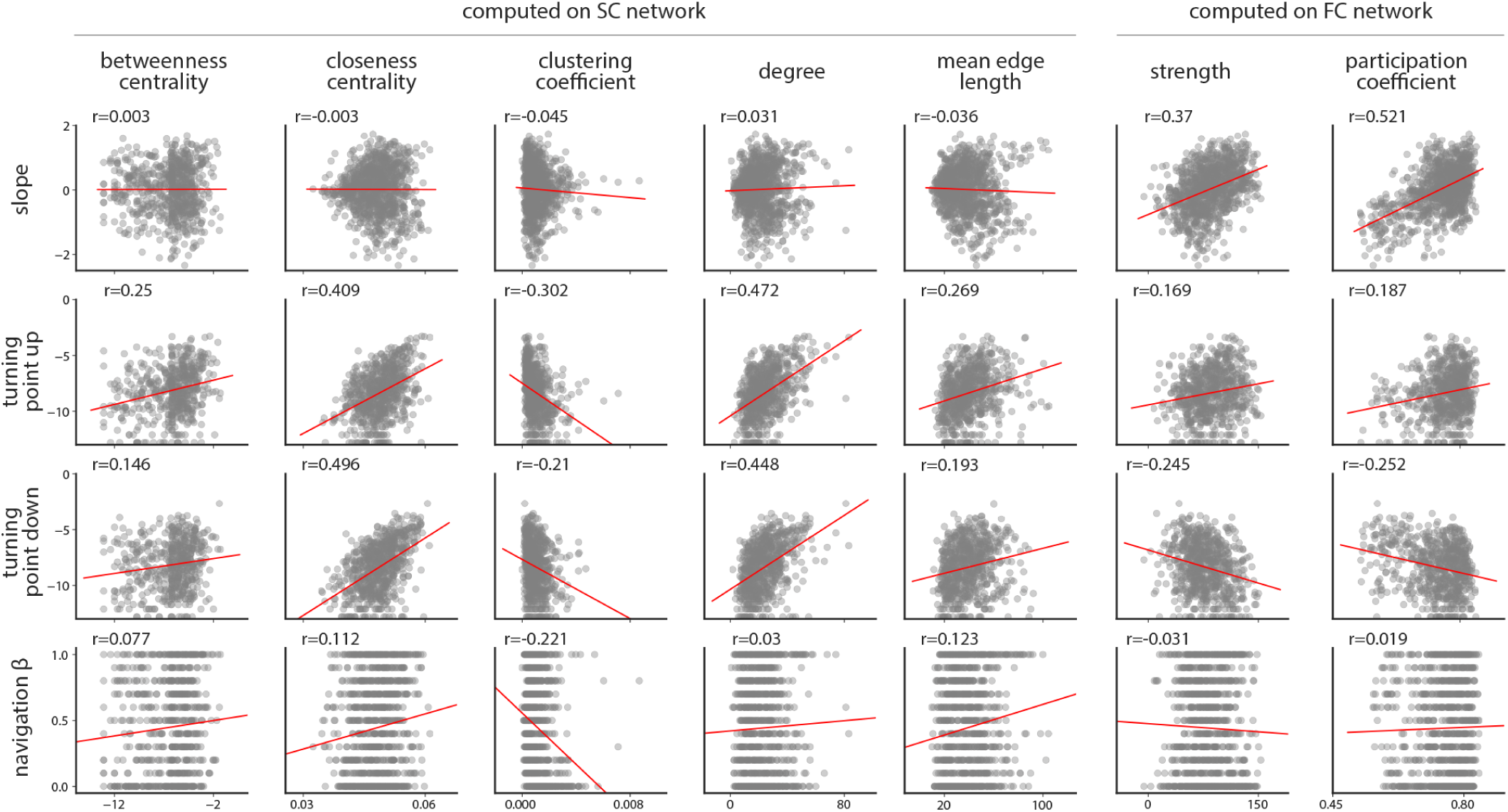
Relating path motif measures with graph properties. The present report derives four node-level dependent variables (mean slope, turning up/down point probability and navigation preference). To assess the extent to which these measures are related to simpler graph properties, we correlate them with node-level features from same structural and functional connectivity matrices. Path motif measures are shown in the rows and graph measures are shown in the columns. The first five graph measures (betweenness, closeness, clustering, degree and mean edge length) are computed on the structural network; the last two graph measures (strength and participation) are computed on the functional network. Edge length refers to physical length and is measured in mm. Participation coefficient is computed with respect to the intrinsic network partition provided by Yeo and colleagues [65]. The two turning point measures and betweenness centrality are log transformed. Relationships are reported in terms of Pearson correlation coefficients.

**Figure S4.**
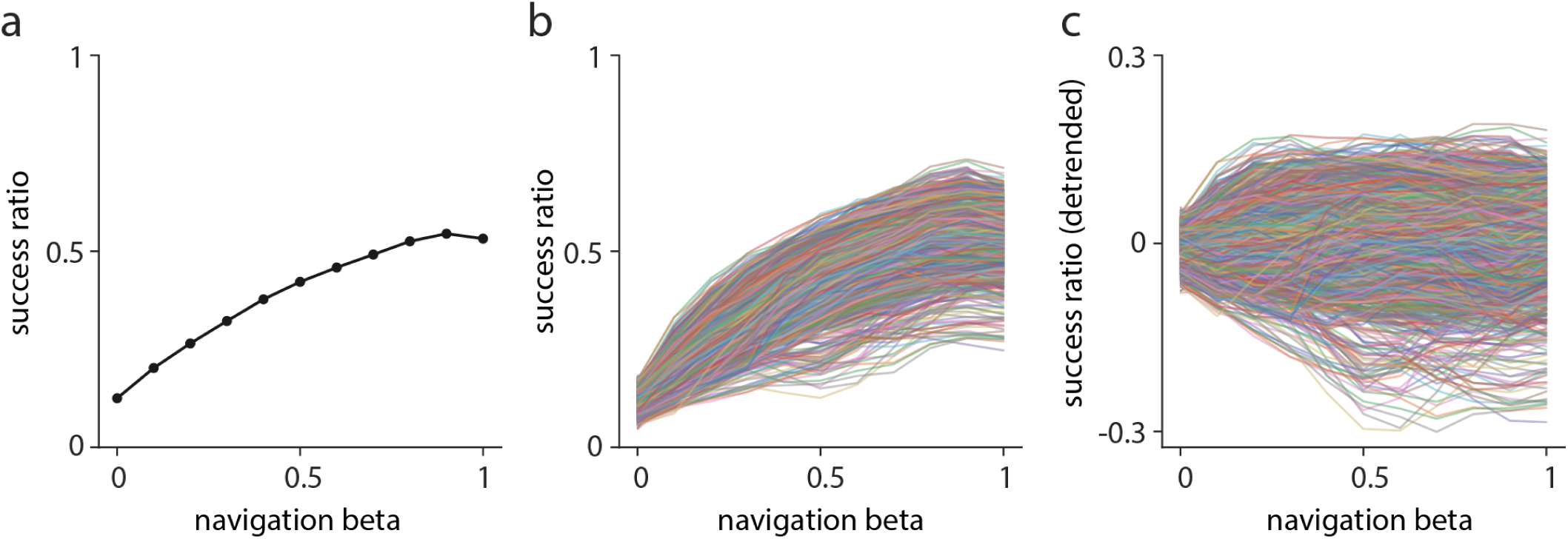
Navigation via space and hierarchy. To assess the use of spatial and hierarchical information to navigate, we derive the proportion of shortest paths successfully recovered (success ratio) as a function *β*, which tunes preference for hierarchy (*β* = 0) vs spatial (*β* = 1.0) information. (a) When the same *β* is imposed on all nodes, there is a stronger preference for spatial navigation, with the optimal *β* = 0.9. (b) Fitting *β* separately for each source node, we observe the same preference for spatial navigation, but also substantial variability across nodes. (c) To ask whether some nodes additionally benefit from hierarchical information, above and beyond spatial information, we detrend the curves and focus on the residual success ratios.

## References

[1] Amico, E., Abbas, K., Duong-Tran, D. A., Tipnis, U., Rajapandian, M., Chumin, E., Ventresca, M., Harezlak, J., and Goñi, J. (2019). Towards a mathematical theory of communication for the human connectome. arXiv preprint 1911.02601.

[2] Avena-Koenigsberger, A., Mišić, B., Hawkins, R. X., Griffa, A., Hagmann, P., Goñi, J., and Sporns, O. (2017). Path ensembles and a tradeoff between communication efficiency and resilience in the human connectome. Brain Struct Funct, 222(1):603–618.

[3] Avena-Koenigsberger, A., Misic, B., and Sporns, O. (2018). Communication dynamics in complex brain networks. Nat Rev Neurosci, 19(1):17–33.

[4] Avena-Koenigsberger, A., Yan, X., Kolchinsky, A., van den Heuvel, M., Hagmann, P., and Sporns, O. (2019). A spectrum of routing strategies for brain networks. PLoS Comput Biol, 15(3):e1006833.

[5] Bertolero, M. A., Yeo, B. T., and D’Esposito, M. (2017). The diverse club. Nat Commun, 8(1):1–11.

[6] Betzel, R. F., Avena-Koenigsberger, A., Goñi, J., He, Y., De Reus, M. A., Griffa, A., Vértes, P. E., Mišic, B., Thiran, J.-P., Hagmann, P., et al. (2016). Generative models of the human connectome. NeuroImage, 124:1054–1064.

[7] Betzel, R. F., Griffa, A., Hagmann, P., and Mišić, (2018a). Distance-dependent consensus thresholds for generating group-representative structural brain networks. Net Neurosci, 3(2):475–496.

[8] Betzel, R. F., Medaglia, J. D., and Bassett, D. S. (2018b). Diversity of meso-scale architecture in human and non-human connectomes. Nature communications, 9(1):346.

[9] Burt, J. B., Demirtaş, M., Eckner, W. J., Navejar, N. M., Ji, J. L., Martin, W. J., Bernacchia, A., Anticevic, A., and Murray, J. D. (2018). Hierarchy of transcriptomic specialization across human cortex captured by structural neuroimaging topography. Nat Neurosci, 21(9):1251.

[10] Cammoun, L., Gigandet, X., Meskaldji, D., Thiran, J. P., Sporns, O., Do, K. Q., Maeder, P., Meuli, R., and Hagmann, P. (2012). Mapping the human connectome at multiple scales with diffusion spectrum mri. J Neurosci Meth, 203(2):386–397.

[11] Coifman, R. R., Lafon, S., Lee, A. B., Maggioni, M., Nadler, B., Warner, F., and Zucker, S. W. (2005). Geometric diffusions as a tool for harmonic analysis and structure definition of data: Diffusion maps. Proc Natl Acad Sci USA, 102(21):7426–7431.

[12] Daducci, A., Gerhard, S., Griffa, A., Lemkaddem, A., Cammoun, L., Gigandet, X., Meuli, R., Hagmann, P., and Thiran, J.-P. (2012). The connectome mapper: an open-source processing pipeline to map connectomes with mri. PLoS ONE, 7(12):e48121.

[13] de Pasquale, F., Della Penna, S., Snyder, A. Z., Marzetti, L., Pizzella, V., Romani, G. L., and Corbetta, M. (2012). A cortical core for dynamic integration of functional networks in the resting human brain. Neuron, 74(4):753–764.

[14] de Reus, M. A. and van den Heuvel, M. P. (2013). Estimating false positives and negatives in brain networks. NeuroImage, 70:402–409.

[15] Demirtaş, M., Burt, J. B., Helmer, M., Ji, J. L., Adkinson, B. D., Glasser, M. F., Van Essen, D. C., Sotiropoulos, S. N., Anticevic, A., and Murray, J. D. (2019). Hierarchical heterogeneity across human cortex shapes large-scale neural dynamics. Neuron, 101(6):1181–1194.

[16] Desikan, R. S., Ségonne, F., Fischl, B., Quinn, B. T., Dickerson, B. C., Blacker, D., Buckner, R. L., Dale, A. M., Maguire, R. P., Hyman, B. T., et al. (2006). An automated labeling system for subdividing the human cerebral cortex on mri scans into gyral based regions of interest. NeuroImage, 31(3):968–980.

[17] Fair, D. A., Dosenbach, N. U., Church, J. A., Cohen, A. L., Brahmbhatt, S., Miezin, F. M., Barch, D. M., Raichle, M. E., Petersen, S. E., and Schlaggar, B. L. (2007). Development of distinct control networks through segregation and integration. Proc Natl Acad Sci USA, 104(33):13507–13512.

[18] Floyd, R. W. (1962). Algorithm 97: shortest path. Communications of the ACM, 5(6):345.

[19] Fulcher, B. D., Murray, J. D., Zerbi, V., and Wang, X.-J. (2019). Multimodal gradients across mouse cortex. Proc Natl Acad Sci USA, 116(10):4689–4695.

[20] Goñi, J., van den Heuvel, M. P., Avena-Koenigsberger, A., de Mendizabal, N. V., Betzel, R. F., Griffa, A., Hagmann, P., Corominas-Murtra, B., Thiran, J.-P., and Sporns, O. (2014). Resting-brain functional connectivity predicted by analytic measures of network communication. Proc Natl Acad Sci USA, 111(2):833–838.

[21] Gollo, L. L., Mirasso, C., Sporns, O., and Breakspear, M. (2014). Mechanisms of zero-lag synchronization in cortical motifs. PLoS Comput Biol, 10(4).

[22] Gollo, L. L., Roberts, J. A., and Cocchi, L. (2017). Mapping how local perturbations influence systems-level brain dynamics. NeuroImage, 160:97–112.

[23] Goulas, A., Zilles, K., and Hilgetag, C. C. (2018). Cortical gradients and laminar projections in mammals. Trends Neurosci, 41(11):775–788.

[24] Graham, D. and Rockmore, D. (2011). The packet switching brain. J Cogn Neurosci, 23(2):267–276.

[25] Griffa, A., Alemán-Gómez, Y., and Hagmann, P. (2019). Structural and functional connectome from 70 young healthy adults [data set]. Zenodo.

[26] Gulyás, A., Bíró, J. J., Kőrösi, A., Rétvári, G., and Krioukov, D. (2015). Navigable networks as nash equilibria of navigation games. Nat Commun, 6:7651.

[27] Hagmann, P., Cammoun, L., Gigandet, X., Meuli, R., Honey, C. J., Wedeen, V. J., and Sporns, O. (2008). Mapping the structural core of human cerebral cortex. PLoS Biol, 6(7).

[28] Hilgetag, C. C. and Kaiser, M. (2004). Clustered organization of cortical connectivity. Neuroinformatics, 2(3):353–360.

[29] Honey, C. J., Thesen, T., Donner, T. H., Silbert, L. J., Carlson, C. E., Devinsky, O., Doyle, W. K., Rubin, N., Heeger, D. J., and Hasson, U. (2012). Slow cortical dynamics and the accumulation of information over long timescales. Neuron, 76(2):423–434.

[30] Huntenburg, J. M., Bazin, P.-L., Goulas, A., Tardif, C. L., Villringer, A., and Margulies, D. S. (2017). A systematic relationship between functional connectivity and intracortical myelin in the human cerebral cortex. Cereb Cortex, 27(2):981–997.

[31] Jones, D., Knösche, T., and Turner, R. (2013). White matter integrity, fiber count, and other fallacies: the do’s and don’ts of diffusion mri. NeuroImage, 73:239–254.

[32] Kaiser, M. and Hilgetag, C. C. (2006). Nonoptimal component placement, but short processing paths, due to long-distance projections in neural systems. PLoS Comput Biol, 2(7):e95.

[33] Kiebel, S. J., Daunizeau, J., and Friston, K. J. (2008). A hierarchy of time-scales and the brain. PLoS Comput Biol, 4(11):e1000209.

[34] Knock, S., McIntosh, A., Sporns, O., Kötter, R., Hagmann, P., and Jirsa, V. (2009). The effects of physiologically plausible connectivity structure on local and global dynamics in large scale brain models. J Neurosci Meth, 183(1):86–94.

[35] Maier-Hein, K. H., Neher, P. F., Houde, J.-C., Côté, M.-A., Garyfallidis, E., Zhong, J., Chamberland, M., Yeh, F.-C., Lin, Y.-C., Ji, Q., et al. (2017). The challenge of mapping the human connectome based on diffusion tractography. Nat Commun, 8(1):1349.

[36] Margulies, D. S., Ghosh, S. S., Goulas, A., Falkiewicz, M., Huntenburg, J. M., Langs, G., Bezgin, G., Eickhoff, S. B., Castellanos, F. X., Petrides, M., Jefferies, E., and Smallwood, J. (2016). Situating the default-mode network along a principal gradient of macroscale cortical organization. Proc Natl Acad Sci USA, 113(44):12574–12579.

[37] Melozzi, F., Bergmann, E., Harris, J. A., Kahn, I., Jirsa, V., and Bernard, C. (2019). Individual structural features constrain the mouse functional connectome. Proc Natl Acad Sci USA, 116(52):26961–26969.

[38] Mesulam, M.-M. (1998). From sensation to cognition. Brain, 121(6):1013–1052.

[39] Mišić, B., Betzel, R. F., Griffa, A., De Reus, M. A., He, Y., Zuo, X.-N., Van Den Heuvel, M. P., Hagmann, P., Sporns, O., and Zatorre, R. J. (2018). Network-based asymmetry of the human auditory system. Cereb Cortex, 28(7):2655–2664.

[40] Mišić, B., Betzel, R. F., Nematzadeh, A., Goñi, J., Griffa, A., Hagmann, P., Flammini, A., Ahn, Y.-Y., and Sporns, O. (2015). Cooperative and competitive spreading dynamics on the human connectome. Neuron, 86(6):1518–1529.

[41] Mišić, B., Goñi, J., Betzel, R. F., Sporns, O., and McIntosh, A. R. (2014a). A network convergence zone in the hippocampus. PLoS Comput Biol, 10(12):e1003982.

[42] Mišić, B., Sporns, O., and McIntosh, A. R. (2014b). Communication efficiency and congestion of signal traffic in large-scale brain networks. PLoS Comput Biol, 10(1):e1003427.

[43] Mohr, H., Wolfensteller, U., Betzel, R. F., Mišić, B., Sporns, O., Richiardi, J., and Ruge, H. (2016). Integration and segregation of large-scale brain networks during short-term task automatization. Nat Commun, 7(1):1–12.

[44] Murray, J. D., Bernacchia, A., Freedman, D. J., Romo, R., Wallis, J. D., Cai, X., Padoa-Schioppa, C., Pasternak, T., Seo, H., Lee, D., et al. (2014). A hierarchy of intrinsic timescales across primate cortex. Nat Neurosci, 17(12):1661.

[45] Muscoloni, A. and Cannistraci, C. V. (2019). Navigability evaluation of complex networks by greedy routing efficiency. Proc Natl Sci Acad USA, 116(5):1468–1469.

[46] Muscoloni, A., Thomas, J. M., Ciucci, S., Bianconi, G., and Cannistraci, C. V. (2017). Machine learning meets complex networks via coalescent embedding in the hyperbolic space. Nat Commun, 8(1):1–19.

[47] Oldham, S., Fulcher, B., Parkes, L., Arnatkeviciūtė, A., Suo, C., and Fornito, A. (2019). Consistency and differences between centrality measures across distinct classes of networks. PLoS ONE, 14(7).

[48] Paquola, C., Wael, R. V. D., Wagstyl, K., Bethlehem, R. A. I., Hong, S.-J., Seidlitz, J., Bullmore, E. T., Evans, A. C., Misic, B., Margulies, D. S., Smallwood, J., and Bernhardt, B. C. (2019). Microstructural and functional gradients are increasingly dissociated in transmodal cortices. PLoS Biol, 17(5):e3000284.

[49] Power, J. D., Barnes, K. A., Snyder, A. Z., Schlaggar, B. L., and Petersen, S. E. (2012). Spurious but systematic correlations in functional connectivity mri networks arise from subject motion. NeuroImage, 59(3):2142–2154.

[50] Roberts, J. A., Perry, A., Roberts, G., Mitchell, P. B., and Breakspear, M. (2017). Consistency-based thresholding of the human connectome. NeuroImage, 145:118–129.

[51] Rosvall, M., Esquivel, A. V., Lancichinetti, A., West, J. D., and Lambiotte, R. (2014). Memory in network flows and its effects on spreading dynamics and community detection. Nat Commun, 5:4630.

[52] Roy, B. (1959). Transitivité et connexité. Comptes Rendus Hebdomadaires Des Seances De L Academie Des Sciences, 249(2):216–218.

[53] Seguin, C., Razi, A., and Zalesky, A. (2019). Inferring neural signalling directionality from undirected structural connectomes. Nat Commun, 10(1):1–13.

[54] Seguin, C., Van Den Heuvel, M. P., and Zalesky, A. (2018). Navigation of brain networks. Proc Natl Acad Sci USA, 115(24):6297–6302.

[55] Shine, J. M., Bissett, P. G., Bell, P. T., Koyejo, O., Balsters, J. H., Gorgolewski, K. J., Moodie, C. A., and Poldrack, R. A. (2016). The dynamics of functional brain networks: integrated network states during cognitive task performance. Neuron, 92(2):544–554.

[56] Suarez, L. E., Markello, R. D., Betzel, R. F., and Misic, B. (2020). Linking structure and function in macroscale brain networks. Trends Cogn Sci.

[57] Thomas, C., Frank, Q. Y., Irfanoglu, M. O., Modi, P., Saleem, K. S., Leopold, D. A., and Pierpaoli, C. (2014). Anatomical accuracy of brain connections derived from diffusion mri tractography is inherently limited. Proc Natl Acad Sci USA, 111(46):16574–16579.

[58] van den Heuvel, M. P., Kahn, R. S., Goñi, J., and Sporns, O. (2012). High-cost, high-capacity backbone for global brain communication. Proc Natl Acad Sci USA, 109(28):11372–11377.

[59] Van Der Maaten, L., Postma, E., and Van den Herik, J. (2009). Dimensionality reduction: a comparative. J Mach Learn Res, 10:66–71.

[60] Vázquez-Rodríguez, B., Suárez, L. E., Markello, R. D., Shafiei, G., Paquola, C., Hagmann, P., van den Heuvel, M. P., Bernhardt, B. C., Spreng, R. N., and Misic, B. (2019). Gradients of structure–function tethering across neocortex. Proc Natl Acad Sci USA, 116(42):21219–21227.

[61] Wagstyl, K., Ronan, L., Goodyer, I. M., and Fletcher, P. C. (2015). Cortical thickness gradients in structural hierarchies. NeuroImage, 111:241–250.

[62] Wang, X.-J. (2020). Macroscopic gradients of synaptic excitation and inhibition in the neocortex. Nat Rev Neurosci, pages 1–10.

[63] Warshall, S. (1962). A theorem on boolean matrices. Journal of the ACM.

[64] Worrell, J. C., Rumschlag, J., Betzel, R. F., Sporns, O., and Mišić, B. (2017). Optimized connectome architecture for sensory-motor integration. Net Neurosci, 1(4):415–430.

[65] Yeo, B., Krienen, F. M., Sepulcre, J., Sabuncu, M. R., Lashkari, D., Hollinshead, M., Roffman, J. L., Smoller, J. W., Zöllei, L., Polimeni, J. R., et al. (2011). The organization of the human cerebral cortex estimated by intrinsic functional connectivity. J Neurophysiol, 106(3):1125–1165.

[66] Zalesky, A., Fornito, A., Cocchi, L., Gollo, L. L., van den Heuvel, M. P., and Breakspear, M. (2016). Connectome sensitivity or specificity: which is more important? NeuroImage, 142:407–420.

[67] Zheng, Y.-Q., Zhang, Y., Yau, Y., Zeighami, Y., Larcher, K., Misic, B., and Dagher, A. (2019). Local vulnerability and global connectivity jointly shape neurodegenerative disease propagation. PLoS Biol, 17(11).

